# Variable transmission efficiency of mammalian origin HPAI D1.1 H5N1 strains in ferrets

**DOI:** 10.64898/2026.05.07.722809

**Authors:** Grace E. Quirk, Michelle N. Vu, Valerie Le Sage, Kaitlyn Bushfield-Thomason, Hanh Dung Nguyen, Seema S. Lakdawala

**Author notes:** These authors had equal contributions, and order was determined by coin flip (Grace E. Quirk, Michelle N. Vu).

## Abstract

Highly pathogenic avian influenza H5N1 2.3.4.4b genotype D1.1 lineage continues to predominate in the United States wild bird population and has spilled over into dairy cattle three independent times. To assess the transmission risk of this sublineage, we performed direct-contact transmission experiments for three distinct D1.1 strains in ferrets. Two of these strains were isolated from humans and one from a lethal cat infection. We found that only one human isolate (A/NV/10/2025) was able to transmit efficiently between ferrets. Compared to the other strains, this isolate harbored the mammalian adaptive PB2 D701N mutation, suggesting this mutation may be critical for D1.1 transmission as opposed to the PB2 E627K substitution present in the lethal cat isolate. Based on these data we conclude that the transmission fitness of D1.1 strains is modest but that special attention should be paid to emergence of adaptation at the PB2 701 position.

## Introduction

Highly pathogenic avian influenza (HPAI) H5N1 clade 2.3.4.4b viruses have been causing notable morbidity and mortality in wild birds since 2021. In early 2024, genotype B3.13 spilled over into dairy cattle and has sustained circulation in the bovine population since then. Towards the end of 2024, a new genotype, D1.1, was detected in wild birds. Recently, D1.1 spilled into dairy cattle at least three independent times, spread to other peridomestic animals including cats, and caused at least 24 human infections in the United States [1–5]. D1.1 has caused more severe infections, including two deaths in humans, compared to B3.13 which predominantly results in conjunctivitis and mild respiratory illness [2,6].

Mammalian adaptive mutations have recurrently evolved in both genotypes and have been prevalent in spillover events to novel hosts. Sequencing D1.1 isolates from various hosts has commonly identified viral strains that contain mammalian adaptations in the PB2 gene. Specifically, D1.1 strains have repeatedly acquired either E627K or D701N, known mammalian adaptive mutations [4], compared to B3.13 sublineage where M631L is fixed [7]. The acquisition of known mammalian PB2 mutations increases the pandemic risk of these viruses as both PB2 627 and 701 have been associated with successful spillover events [7].

In this study, we examined the direct contact transmission fitness of three mammalian origin D1.1 strains (**Figure 1 and Table S1**).

**Figure 1.**
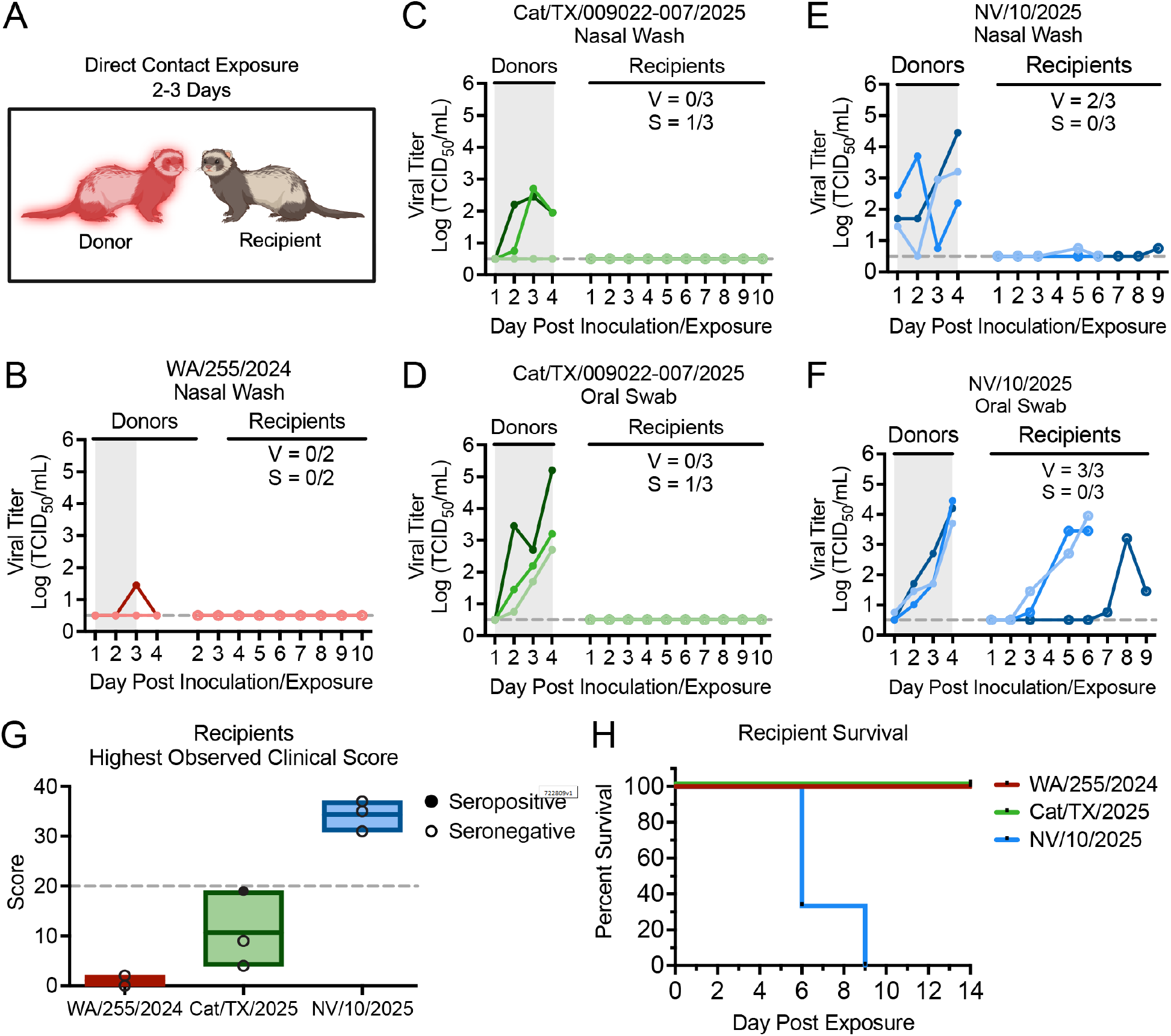
D1.1 A/NV/10/2025 human isolate efficiently transmits in ferrets by direct contact exposure. **(A)** Recipient ferrets with no prior immunity were exposed to inoculated donor ferrets for 2-3 days by direct contact, starting from day 1 post donor inoculation. All donor ferrets were inoculated intranasally with 10^4^-10^5^ TCID_50_/500µL of each specified virus. **(B-F)** Viral shedding by both donors and recipients was measured in nasal washes and oral swabs by TCID_50_ assay for D1.1 strains A/WA/255/2024 **(B)**, A/Cat/TX/009022-007/2025 **(C, D)** and A/NV/10/2025 **(E, F)**. Exposure window is indicated in grey. The number of recipients that shed virus (V) or seroconverted (S) are indicated in the fractions. **(G)** The highest observed clinical sign score for recipients over the course of the study is shown. Ferrets reached humane endpoint with a clinical score at or above 20 (dotted line). **(H)** Recipient percent survival is shown for each strain.

## Results

To assess the pandemic risk of this genotype, we evaluated the transmissibility of three D1.1 strains of human or domestic cat origin (**Table S1**) between ferrets by direct contact. The three chosen strains contained several other mutations across all eight gene segments (**Table S2**) but were selected as representative strains with distinct PB2 mammalian adaptive mutations. To assess transmissibility, we inoculated donor ferrets intranasally with either A/WA/255/2024 (PB2 627E and 701D), A/Cat/TX/009022-007/2025 (PB2 627K and 701D), or A/NV/10/2025 (PB2 627E and 701N). Donors were cohoused with immunologically naïve recipients for 2-3 days starting from day 1 post donor inoculation before being separated and single-housed for the remainder of the study (**Fig 1A**). Donors were euthanized at predetermined time points post exposure but before humane endpoints were reached, while recipients were monitored until 14 days post exposure or until humane endpoints were reached.

We first assessed A/WA/255/2024, a strain harboring avian residues PB2 627E and 701D isolated from a mild human infection due to commercial poultry exposure (**Table S1)**. Only one of the two donor ferrets intranasally inoculated with 10^4^ TCID_50_ of A/WA/255/2024 shed virus in nasal washes at low levels during the exposure window (**Fig 1B**). Donor ferrets presented with increased temperature within the first 4 days, but otherwise mild clinical signs (**Table S3**). Recipient ferrets which were exposed for 2 days, remained asymptomatic and no viral shedding was detected during the duration of the experiment (**Fig 1B, Fig 1G, and Table S3**). Both recipients were confirmed seronegative by microneutralization with serum collected 14 days post exposure (**Table S3**), indicating a lack of direct-contact transmission of A/WA/255/2024. Based on this initial study, we elected to use a higher viral dose (10^5^ TCID_50_) to inoculate three donor ferrets, a 3-day exposure window, and inclusion of oral swabs to assess viral replication for the remaining D1.1 strains.

A/Cat/TX/009022-007/2025, a lethal cat isolate from exposure to an unknown source, was selected for evaluation based on the presence of mammalian adaptive PB2 E627K mutation (**Table S1**). The three intranasally inoculated donors displayed weight loss and elevated temperatures, and one exhibited hypothermia during the first 4 days of infection (**Table S3**). Two of three donors shed infectious virus in nasal washes while all three donors shed virus in oral swabs during the exposure window (**Fig 1C and 1D**). Despite this robust shedding, no recipient ferrets shed detectable virus in their nasal or oral cavities (**Fig 1C and 1D**). Two recipients showed mild clinical signs, while one displayed more moderate clinical signs including weight loss >20% and rapid respiratory rate (**Fig 1G and Table S3**). This animal recovered prior to reaching endpoint criteria such that all recipient animals survived to day 14 (**Fig 1H**). Sera collected from recipients at day 14 post exposure revealed that the one recipient with more severe clinical signs had detectable anti-H5 antibodies measured by ELISA (**Table S3**). These results demonstrate that although this strain replicated efficiently in donors, it transmitted inefficiently (1/3, 33%) by direct contact.

Lastly, we assessed A/NV/10/2025, a human isolate that harbors the mammalian adaptive PB2 701N residue (**Table S1**). This strain was isolated from a person presenting with mild to moderate symptoms and known exposure to dairy cattle (**Table S1**) [8]. All three inoculated donor ferrets shed infectious virus in nasal washes and oral swabs (**Fig 1E and 1F**). Donor ferrets exhibited severe signs of disease within the first 4 days post inoculation (**Table S3**). Importantly, two recipients shed infectious virus in nasal washes (**Fig 1E**), and all three shed high levels of virus in their oral swabs (**Fig 1F**). Recipients displayed similar clinical signs as donors (**Fig 1G and Table S3)** before meeting humane endpoint at days 6 or 9 post exposure (**Fig 1H**). These results indicate robust (3/3, 100%) direct contact transmission of A/NV/10/2025 in ferrets.

## Discussion

In this study we describe transmission fitness of three D1.1 genotype 2.3.4.4b clade H5N1 viruses. Two of the three strains, A/WA/255/2024 and A/Cat/TX/009022-007/2025, had low to no spread to cohoused ferrets. Reports by the CDC using a related D1.1 strain A/WA/239/2024 reported low transmission efficiency by direct contact to 1/3 recipient ferrets [9]. In contrast, A/NV/10/2025 spread to all three cohoused animals. Additionally, recipient ferrets naturally infected with A/NV/10/2025 reached humane endpoint criteria prior to the scheduled end of the study, suggesting that not only is this virus transmissible, but it can result in severe natural infection.

The viruses in this study were chosen because of their mammalian origin and their unique PB2 mammalian adaptive residues. A/NV/10/2025 has a classical PB2 mammalian adaptive mutation at position 701, in contrast to the 627 residue present in the A/Cat/TX/009022-007/2025 strain. While we cannot exclude the impact of other mutations in present in A/NV/10/2025 (**Table S2**) compared to A/Cat/TX/009022-007/2025 and A/WA/255/2024, it is highly likely that mammalian adaptation of residue 701 is playing a role in the forward transmission potential of this strain. Interestingly, only 1% of all D1.1 strains curated by Nextstrain [10] contain a PB2 D701N mutation, but 7% of spillovers into mammalian species contain this mutation [3]. While PB2 E627K mutation is more prevalent in mammalian spillover viruses at 35%, this mammalian adaptation in D1.1 has not been related to forward transmission of the virus in ferrets in our study or others [9].

Our findings suggest that while less common in initial mammalian spillover events, PB2 D701N may increase the risk of onward transmission. Targeted surveillance of D1.1 strains for the acquisition of PB2 D701N in animals at the human-animal interface may identify priority variants for pandemic risk assessment.

## Methods

Detailed methods can be found in the Supplementary Material.

## Acknowledgements

We thank members of the Lakdawala lab for thoughtful feedback and discussion. We thank Dr. Rachel Duron for editing assistance. This project has been funded in part with Federal funds from the National Institute of Allergy and Infectious Diseases (NIAID), National Institutes of Health (NIH), Department of Health and Human Services, under Contract No. 75N93021C00015 and 75N93021C00017. Operations of The University of Pittsburgh Regional Biocontainment Laboratory (RBL) within the Center for Vaccine Research (CVR) are supported by an NIH award (UC7AI180311) from NIAID.

## List of Supplementary Material

**Table S1.**
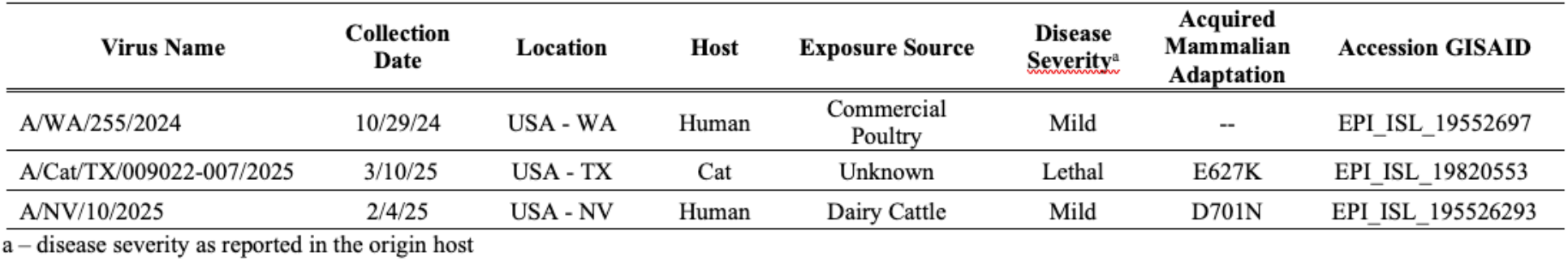
Summary of origin for selected D1.1 H5N1 strains.

**Table S2.**
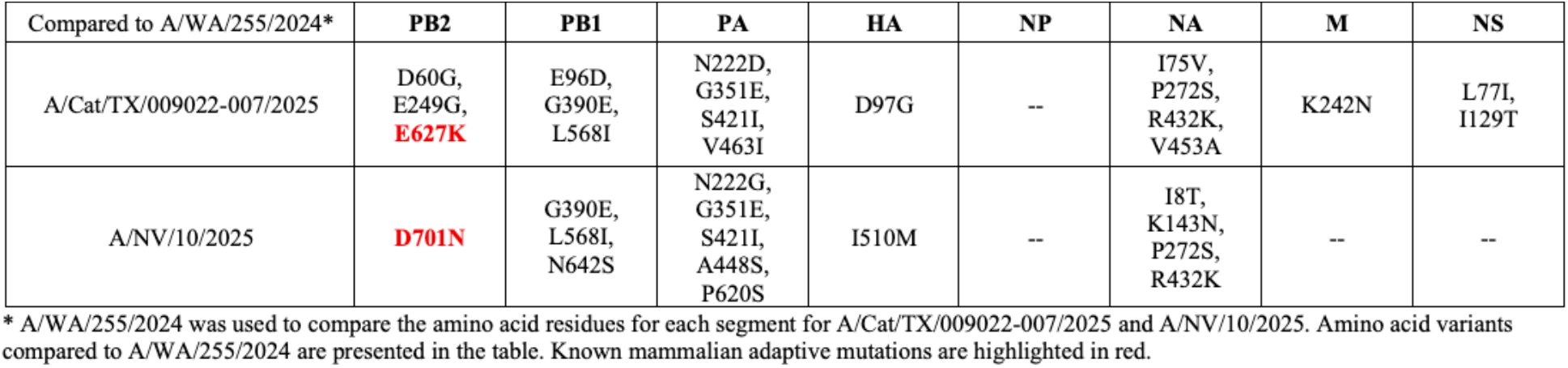
Comparison of altered residues in all eight genes segments between three selected strains.

**Table S3.**
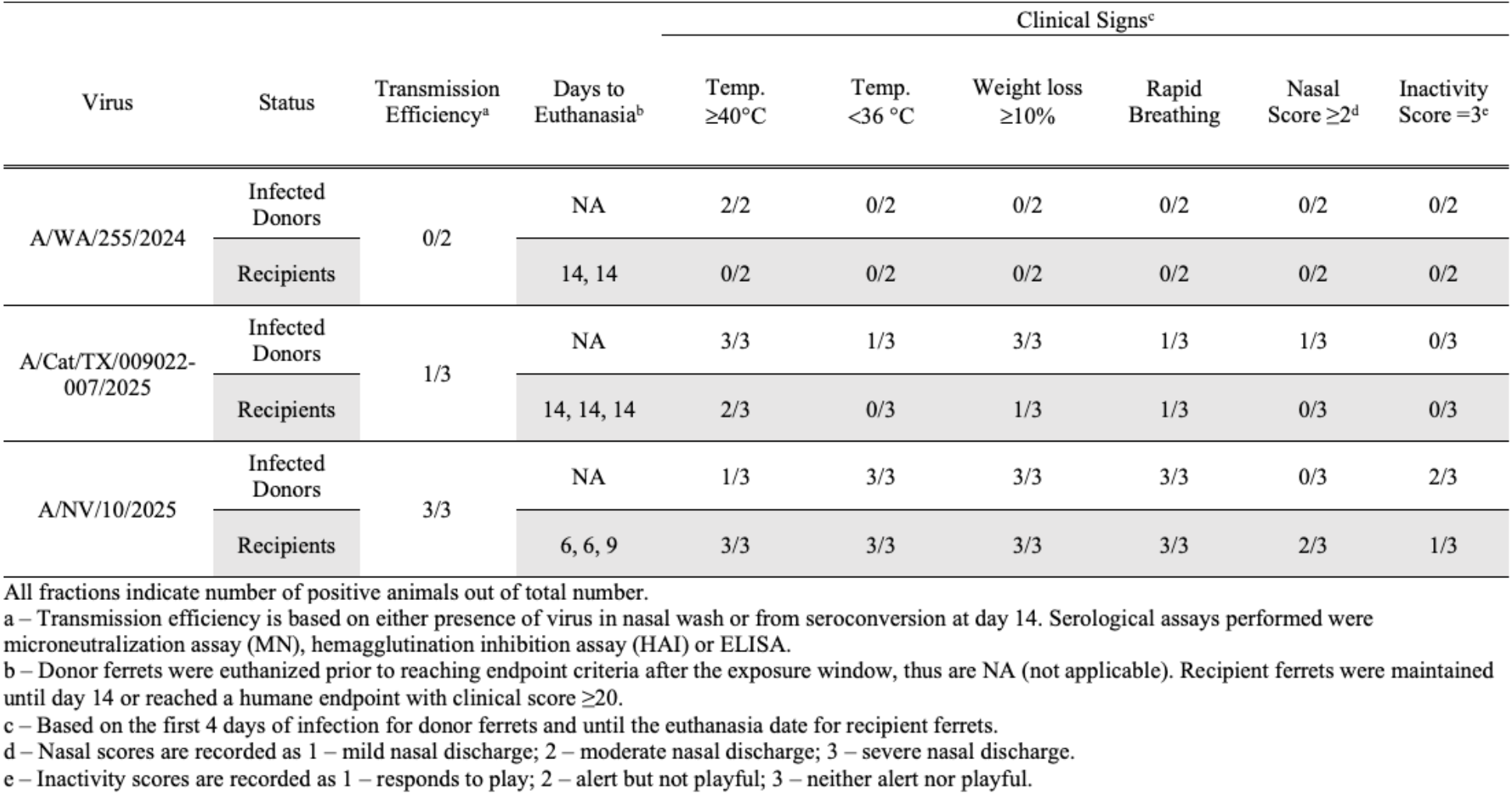
Summary of transmission and clinical signs for Dl.l infected donors and recipients.

## Methods

### Ethics statement

All ferret experiments were conducted in the University of Pittsburgh’s or Emory University’s BSL3 facility in compliance with the guidelines of the Institutional Animal Care and Use Committee (University of Pittsburgh’s approved protocol 24125091; Emory University’s approved protocol 2024-00000112). Isoflurane was used to sedate animals for all nasal washes and oral swabs, as directed by approved methods. For terminal procedures, animals received ketamine and xylazine for sedation, followed by euthanasia solution administered via cardiac injection.

### Cells

Madin-Darby canine kidney (MDCK) and 293T cells were obtained from American Type Culture Collection (ATCC). MDCK cells were maintained in Minimum Essential Medium (MEM) supplemented with 10% fetal bovine serum, 2 mM L-glutamine and 1x penicillin (100 U/mL) and streptomycin (100 µg/mL). 293T cells were maintained in Dulbecco’s Modified Eagle Medium (DMEM) high glucose (4.5 g/L) supplemented with 10% fetal bovine serum and 1x penicillin (100 U/mL) and streptomycin (100 µg/mL). Cells were incubated at 37°C with 5% CO_2_

### Virus Rescue

Reverse genetics plasmids expressing A/WA/255/2024, A/Cat/TX/009022-007/2025, and A/NV/10/2025 were synthesized based on sequence deposited in GISAID (Accession numbers EPI_ISL_19552697, EPI_ISL_19820553, and EPI_ISL_195526293, respectively). Noncoding regions (NCRs) were based on a set of pan-avian H5Nx assembled trees to identify the most conserved sequences across H5Nx strains. The eight reverse genetics plasmids were transfected into 293T cells using Lipofectamine 2000 in Opti-MEM complete media. Twenty-four hours later, supernatant from transfected 293T cells were overlayed on top of MDCK cells with MEM supplemented with 4 mM L-glutamine, 1x antibiotic-antimycotic and 1 µg/mL of TPCK-treated trypsin. The 293T supernatant overlay on top of MDCK cells was repeated at 48 hours post transfection. MDCK cells were monitored for cytopathic effect (CPE) for 2 days post final supernatant overlay, and supernatant was collected (cell passage 1, cP1). A second cell passage (cP2) stock was generated on MDCK cells and used for all subsequent experiments. Plasmids were sequence verified prior to rescue.

### Ferret Screening

Prior to purchase from Triple F Farms (Sayre, PA), sera from six-to eight-month-old male ferrets were screened for antibodies against influenza A and B viruses using hemagglutination inhibition (HAI). Antigens used for screening were obtained through the International Reagent Resource, Influenza Division, WHO Collaborating Center for Surveillance, Epidemiology and Control of Influenza, Centers for Disease Control and Prevention, Atlanta, GA, USA: CDC Antigen, Influenza A(H1N1)pdm09 Control Antigen (A/Victoria/2570/2019), BPL-Inactivated, FR-1820; CDC Antigen, Influenza A(H3) Control Antigen (A/Delaware/01/2021), BPL-Inactivated, FR-1821; CDC Antigen, Influenza B Control Antigen, Victoria Lineage (B/Michigan/01/2021), BPL-Inactivated, FR-1822.

### Transmission Study

Donor ferrets were intranasally inoculated with 10^4^ TCID_50_ (for A/WA/255/2024) or 10^5^ TCID_50_ (for A/Cat/TX/009022-007/2025 or A/NV/10/2025) in 500 µl. At day 1 post donor inoculation, donors and recipients were paired and cohoused for 2 (for A/WA/255/2024) or 3 days (for A/Cat/TX/009022-007/2025 or A/NV/10/2025) of direct-contact exposure. Pairs were separated and single-housed after exposure. Nasal washes and oral swabs were collected from donors for 4 days post inoculation before euthanasia at predetermined time points. Nasal washes and oral swabs were collected from recipients for 10 days post exposure or until reaching humane endpoint criteria. Sera was collected from recipients at 14 days post exposure or upon reaching humane endpoint and used for serological assays described below. Recipients were euthanized at 14 days post exposure if not euthanized early due to reaching humane endpoint criteria.

### Ferret Clinical Signs

Ferrets were monitored for signs of clinical disease throughout the study to assess disease severity with scoring expanded upon what was previously described [1]. Humane endpoint is met at a cumulative clinical score ≥20. Clinical signs were assigned point values as follows: Activity (0 = alert and playful; 2 = alert but playful only when stimulated; 4 = alert but not playful when stimulated; 10 = neither alert nor playful when stimulated; 20 = moribund), Temperature (0 = normal 36-40°C; 2 = fever ≥40°C; 4 = severe fever ≥40°C + 2°C more than baseline; 20 = hypothermia <36°C), Weight Loss (0 = 0-5%; 2 = 5-10%; 4 = 10-15%; 6 = 15-20%; 10 = 20-22%; 20 = ≥22%), Nasal Discharge (0 = none; 1 = mild; 2 = moderate; 3 = severe), Respiratory Rate (0 = normal breathing; 5 = rapid breathing), Dehydration (0 = none; 3 = skin tent ≥1 seconds), and Diarrhea (0 = none; 1 = present).

### Viral Titration

For TCID_50_ virus titration, samples were titrated in both 24-well and 96-well plates. For 24-well plates, 100 µl of samples were added in duplicate to confluent MDCK cells in MEM supplemented with 4 mM L-glutamine, 1x antibiotic-antimycotic and 1 µg/mL TPCK-treated trypsin. For 96-well plates, ten-fold serial dilutions were made with samples in MEM supplemented with 4 mM L-glutamine, 1x antibiotic-antimycotic and 1 µg/mL TPCK-treated trypsin and added to confluent MDCK cells in quadruplicate. MDCK cells were observed after 4 days for cytopathic effect (CPE). Virus titers were calculated using Reed and Muench method [2] and expressed as log_10_ TCID_50_/mL. The limit of detection for this assay is 10^0.5^ TCID_50_/mL.

### Microneutralization Assay

Sera was RDE-treated with 3 volumes of 2X RDE to 1 volume of sera overnight at 37°C. Sera was then heat inactivated at 56°C for 30 minutes and 6 volumes of saline was added for a final dilution of 1:10. The titer of neutralizing antibodies was determined by two-fold serial dilutions of RDE-treated ferret serum in a volume of 125 µL was incubated with 125 µL of 10^3.3^ TCID_50_ of virus for 1 hour at room temperature with continuous rocking. 150 µL of MEM supplemented with 2 mM L-glutamine and 1X antibiotic-antimycotic with TPCK was added to 96-well plates with confluent MDCKs before 50 µL of virus:serum mixture was added in quadruplicate. After 4 days, CPE was determined, and the neutralizing antibody titer was expressed as the reciprocal of the highest dilution of serum required to completely neutralize the infectivity of each virus on MDCK cells. The concentration of antibody required to neutralize 100 TCID_50_ of virus was calculated based on the neutralizing titer dilution divided by the initial dilution factor, multiplied by the antibody concentration. The limit of detection for this assay is 14.

### HAI

RDE-treated sera were serially diluted two-fold and incubated with eight hemagglutinating units of HA antigen or virus. After 15 minutes of incubation, an equal volume of 0.75% turkey red blood cells (Lampire Biological Laboratories) was added and incubated for 30 minutes. The reciprocal of the highest dilution of serum that inhibited hemagglutination was determined to be the HAI titer. The limit of detection for this assay is 1:10.

### ELISA

Influenza A H5N1 clade 2.3.4.4b (A/Louisiana/12/2024) Hemagglutinin protein was purchased from Sino Biological (cat: 41060-V08B). Antigens were immobilized on high-capacity binding 96 well-plates (Corning, 9018) overnight in sterile phosphate-buffered saline (PBS) at 4 ng/mL. Coated plates were washed with a PBS containing 0.05% Tween-20 (PBS-T) buffer. Plates were blocked with 5% non-fat milk (ChemCruz Botto Dry Milk, sc2325) in PBS-T for 1 hour. Blocking solution was removed and plates were washed with PBS-T. Ferret serum was initially diluted 1:40 in PBS and serially diluted in duplicate with seven 1:3 dilutions and overlaid onto the coated plate for 1 hour. Plates were washed and incubated with 5% non-fat milk in PBS-T containing a 1:2000 dilution of Goat Anti-Ferret IgG H&L (HRP) (Abcam, ab112770) for 1 hour in the dark. Plates were washed with PBS-T and then PBS. Plates were developed in 1-Step TMB Substrate (Thermo Fisher Scientific) and quenched with 2N H_2_SO_4_. Plates were read at 450nm with a SpectraMax 340PC384 Microplate Reader (Molecular Devices). Area under the curve (AUC) was calculated using GraphPad Prism (v10).

