## Supplemental Table S1 for "Variable transmission efficiency of mammalian origin HPAI D1.1 H5N1 strains in ferrets"

**Table S1: Summary of origin for selected D1.1 H5N1 strains**

| Virus Name | Collection Date | Location | Host | Exposure Source | Disease Severity[a] | Acquired Mammalian Adaptation | Accession GISAID |
|---|---|---|---|---|---|---|---|
| A/WA/255/2024 | 10/29/24 | USA - WA | Human | Commercial Poultry | Mild | -- | EPI_ISL_19552697 |
| A/Cat/TX/009022-007/2025 | 3/10/25 | USA - TX | Cat | Unknown | Lethal | E627K | EPI_ISL_19820553 |
| A/NV/10/2025 | 2/4/25 | USA - NV | Human | Dairy Cattle | Mild | D701N | EPI_ISL_195526293 |

a – disease severity as reported in the origin host
