## Supplemental Table S2 for "Variable transmission efficiency of mammalian origin HPAI D1.1 H5N1 strains in ferrets"

**Table S2: Comparison of altered residues in all eight genes segments between three selected strains**

| Compared to A/WA/255/2024* | **PB2** | **PB1** | **PA** | **HA** | **NP** | **NA** | **M** | **NS** |
|---|---|---|---|---|---|---|---|---|
| A/Cat/TX/009022-007/2025 | D60G, E249G, **E627K** | E96D, G390E, L568I | N222D, G351E, S421I, V463I | D97G | -- | I75V, P272S, R432K, V453A | K242N | L77I, I129T |
| A/NV/10/2025 | **D701N** | G390E, L568I, N642S | N222G, G351E, S421I, A448S, P620S | I510M | -- | I8T, K143N, P272S, R432K | -- | -- |

* A/WA/255/2024 was used to compare the amino acid residues for each segment for A/Cat/TX/009022-007/2025 and A/NV/10/2025. Amino acid variants compared to A/WA/255/2024 are presented in the table. Known mammalian adaptive mutations are highlighted in red.
