## Supplemental Table S3 for "Variable transmission efficiency of mammalian origin HPAI D1.1 H5N1 strains in ferrets"

**Table S3: Summary of transmission and clinical signs for D1.1 infected donors and recipients**

| Virus | Status | Transmission Efficiency[a] | Days to Euthanasia[b] | Clinical Signs[c] | | | | | |
|---|---|---|---|---|---|---|---|---|---|
| | | | | Temp. ≥40°C | Temp. <36 °C | Weight loss ≥10% | Rapid Breathing | Nasal Score ≥2[d] | Inactivity Score =3[e] |
| A/WA/255/2024 | Infected Donors | 0/2 | NA | 2/2 | 0/2 | 0/2 | 0/2 | 0/2 | 0/2 |
| | Recipients | | 14, 14 | 0/2 | 0/2 | 0/2 | 0/2 | 0/2 | 0/2 |
| A/Cat/TX/009022-007/2025 | Infected Donors | 1/3 | NA | 3/3 | 1/3 | 3/3 | 1/3 | 1/3 | 0/3 |
| | Recipients | | 14, 14, 14 | 2/3 | 0/3 | 1/3 | 1/3 | 0/3 | 0/3 |
| A/NV/10/2025 | Infected Donors | 3/3 | NA | 1/3 | 3/3 | 3/3 | 3/3 | 0/3 | 2/3 |
| | Recipients | | 6, 6, 9 | 3/3 | 3/3 | 3/3 | 3/3 | 2/3 | 1/3 |

All fractions indicate number of positive animals out of total number.
a – Transmission efficiency is based on either presence of virus in nasal wash or from seroconversion at day 14. Serological assays performed were microneutralization assay (MN), hemagglutination inhibition assay (HAI) or ELISA.
b – Donor ferrets were euthanized prior to reaching endpoint criteria after the exposure window, thus are NA (not applicable). Recipient ferrets were maintained until day 14 or reached a humane endpoint with clinical score ≥20.
c – Based on the first 4 days of infection for donor ferrets and until the euthanasia date for recipient ferrets.
d – Nasal scores are recorded as 1 – mild nasal discharge; 2 – moderate nasal discharge; 3 – severe nasal discharge.
e – Inactivity scores are recorded as 1 – responds to play; 2 – alert but not playful; 3 – neither alert nor playful.
